# Classification of Neurons in the Adult Mouse Cochlear Nucleus: Linear Discriminant Analysis

**DOI:** 10.1101/594713

**Authors:** Paul B. Manis, Michael R. Kasten, Ruili Xie

## Abstract

The cochlear nucleus (CN) transforms the spike trains of spiral ganglion cells into a new set of sensory representations that are essential for auditory discriminations and perception. These transformations require the coordinated activity of different classes of neurons that are embryologically derived from distinct sets of precursors. Decades of investigation have shown that the neurons of the CN are differentiated by their ion channel expression and intrinsic excitability. In the present study we have used linear discriminant analysis (LDA) to perform an unbiased analysis of measures of the responses of CN neurons to current injections to mathematically separate cells on the basis of both morphology and physiology. Recordings were made from cells in brain slices from CBA mice and a transgenic mouse line, NF107, crossed against the Ai32 line. For each cell, responses to current injections were analyzed for spike rate, spike shape (action potential height, afterhyperpolarization depth, first spike half-width), input resistance, resting membrane potential, membrane time constant, hyperpolarization-activated sag and time constant. Cells were filled with dye for morphological classification, and visually classified according to published accounts. The different morphological classes of cells were separated with the LDA. Ventral cochlear nucleus (VCN) bushy cells, planar multipolar (T-stellate) cells, and radiate multipolar (D-stellate) cells were in separate clusters, and were also separated from all of the neurons from the dorsal cochlear nucleus (DCN). Within the DCN, the pyramidal cells and tuberculoventral cells were largely separated from a distinct clusters of cartwheel cells. DCN cells fell largely within a plane in the first 3 principal axes, whereas VCN cells were in 3 clouds approximately orthogonal to this plane. VCN neurons from the two mouse strains were slightly separated, indicating either a strain dependence or the differences in slice preparation methods. We conclude that cochlear nucleus neurons can be objectively distinguished based on their intrinsic electrical properties, but that such distinctions are still best aided by morphological identification.

## Introduction

Neurons of the mammalian cochlear nucleus exhibit a variety of responses to intracellular current injection, reflecting the distinct expression of collections of ion channels amongst different classes. However, even within a class, such as bushy cells, individual cells may express specific conductances at different magnitudes [1–3], leading to diversity in excitability features such as action potential threshold, action potential height, and rheobase. In spite of this variability, cells of a given morphological class appear to possess common properties that have been used to identify cells on the basis of their electrical signatures alone [4–12].

Quantitative methods for identifying cell classes have been explored in the context of the myriad interneuronal populations in cortex [13], within the olfactory bulb [14] and across neuronal populations throughout the brain [15]. These methods rely on systematic measurement of distinct features of intrinsic excitability such as action potential shape, firing rates, passive membrane measures, and responses to hyperpolarization, and have used principal components analysis (PCA), support vector machine model, or stepwise linear regressions. Within the cochlear nucleus, application of an hierarchical clustering analysis to *in vivo* single unit data provided evidence for partial separation of unit response types in the gerbil AVCN [16], although further analysis (using PCA) suggested that there was extensive overlap between cell classes.

Here we apply linear discriminant analysis [17] to the problem of separating cell classes in the cochlear nucleus based on intrinsic excitability. Whereas PCA separates classes by finding the axes that maximize the variance within a data set, and does not rely on labels, LDA maximizes the separation between classes, utilizing label (e.g., class) information. We find that LDA is an effective tool for segregating the cell classes based on their excitability, while also suggesting that there is either overlap between the properties of some of the classes, or that they may not be entirely morphologically distinguishable. Such a classification tool should be useful in future studies of the excitability of cochlear nucleus neurons following hearing loss as a way of objectively assessing how the excitability of neurons changes.

## Materials and Methods

Whole cell tight-seal recordings were made in brain slices from adult CBA (P28-69) and NF107::Ai32 (P31-166) mice. The NF107::Ai32 mice are the F1 cross of the NF107 mouse line, originally from the GENSAT Consosrtium [18], and the channel rhodopsin (ChR2) expressing line Ai32 [19], and so are on a mixed CD-1, C57Bl/6J and FVB background. The ChR2 was not activated during these experiments. CBA mice were of either sex, whereas the NF107::AI32 mice were only males, as the Cre driver is carried on the Y chromosome.The data from the CBA mice were taken from a previous series of studies [11,20]. Data from the NF107::Ai32 mice were taken from unpublished work (Kasten, Ropp and Manis, in preparation). The CBA slices were prepared following anesthesia (100 mg/kg ketamine and 10 mg/kg xylazine), and decapitation, with slicing in warm ACSF. The NF107::Ai32 slices were prepared using the same anesthesia followed by transcardial perfusion with an NMDG-based solution [21]. Electrodes contained 126 K-gluconate, 6 KCl, 2 NaCl, 10 HEPES, 0.2 EGTA, 4 Mg-ATP, 0.3 Tris-GTP, and 10 Tris-phosphocreatine, with pH adjusted to 7.2 with KOH and recordings were made with a MultiClamp 700B (Molecular Devices) amplifier, low-pass filtered at 6kHz, and digitized at 10-20 kHz with 16-bit A-D converters (National Instruments). Stimulus presentation and acquisition were controlled by either a custom Matlab® program or by *acq4* [22]. All animal procedures were approved by the University of North Carolina Institutuional and Animal Care Committee (protocols 12-253, 15-253 and 18-160).

For each cell, responses to current injections (100-500 msec duration, ranging from −1 to 4 nA) were analyzed. Data from either acquisition program were converted to a common format for analysis by Python (V3.6) scripts. Passive measures included input resistance (from the slope of the current-voltage relationship just below rest), resting membrane potential, membrane time constant (measured from responses to small hyperpolarizing current steps that produced 2-10 mV voltage deflections), the magnitude of the hyperpolarization sag [23] and the time constant for the sag measured near −80 mV. Active measures included action potential height (measured from rest to action potential peak), first spike half-width (measured at half the action potential height from rest), afterhyperpolarization depth (measured from rest to the first afterhyperpolarization), an adaptation index measured near firing threshold (see below), the number of rebound spikes after hyperpolarizing steps, the coefficient of variation of interspike intervals, and the slope of the firing rate versus current curve for the first 3 current levels above threshold. Cells were filled with dyes (AlexaFluor 488 for CBA mice; tetramethylrhodamine biocytin for the NF107:Ai32 mice) for morphological classification, and visually classified according to published accounts, based on digital images and image stacks collected at low (4X) and high (40-63X) power either during or immediately after each cell was recorded.

Adaptation was measured for the lowest two levels of current that elicited spikes as:

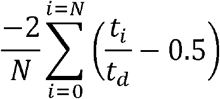

Where *t*_*i*_ is the time of the *i*^*th*^ spike in the trace, *t*_*d*_ is the trace duration, and *N* is the number of spikes. This measure ranges from −1 to 1. Neurons that fire regularly without adaptation throughout the trace will have an index of 0. Neurons that fire preferentially only at the onset of the trace will have an index of 1, whereas those that fire near the end of the trace will have an index of −1. Thus, bushy cells will have an index of 1, stellate cells and tuberculoventral cells will usually have an index near 0, and pyramidal cells may have a negative index, depending on the delay to the first spike. Note that this measure depends on the current level that is used relative to the spike threshold, as well as the current duration. The adaptation measured at the threshold current was found to be uninformative in preliminary analyses, and so the only adaptation computed from the next higher current that evoked spikes was used.

All absolute voltage measurements are corrected for a −11 mV junction potential for the K-gluconate electrodes. All other voltage measurements are differential (action potential height from peak to the minimum of the following AHP) and are independent of the junction potential.

Computed measures were then analyzed using LDA and PCA using standard libraries in Python (scikit-learn v0.20, Python 3.6), and in R (3.5, using the packages DisplayR and flipMultivariates).

## Results

The discharge patterns of cochlear nucleus neurons have been reported in a series of studies over the years from multiple laboratories using similar, but not identical recording conditions. Fig 1 shows the intrinsic physiology of example cells from six major morphological classes as recorded in our dataset. Briefly, bushy cells (Fig 1A) fire 1-3 action potentials at the onset of depolarizing current injections, and are silent thereafter [4,5]. At higher current levels, oscillatory membrane responses, which may represent axonally initiated action potentials, are sometimes visible. In response to hyperpolarizing pulses, bushy cells can show a slow sag in membrane potential, and following the hyperpolarizing step can generate an anodal break spike. The planar multipolar cells (Fig 1B) fire regularly in response to depolarizing current injections, and also show a slow sag in response to hyperpolarizing current steps; they can also show anodal break spikes [11,23]. Radiate multipolar cells also fire regularly in response to depolarization, sometimes exhibiting an adapting spike train. They show a rapid sag in response to hyperpolarization, and frequently have anodal break spikes. As noted previously [11], radiate multipolar cells also may fire only at the onset of a weak depolarizing current pulse. Pyramidal (fusiform) cells of the dorsal cochlear nucleus fire regularly [9,24,25], and may have a long delay to the first spike or a long first interspike interval [25–27]. In the adult mice studies here, these cells do not show a prominent sag, but do show a rapidly activating rectification in response to hyperpolarizing steps, which is likely generated by Kir channels [28]. Cartwheel cells show mixed single-spiking regular firing and burst firing [7,10]. The tuberculoventral neurons show regular firing, and often have trains of re8ound spikes after hyperpolarization [12,29]. The principal cell database included 18 bushy cells, 31 planar multipolar cells, 32 radiate multipolar cells, 38 pyramidal cells, 12 cartwheel cells, and 31 tuberculoventral cells. Additional cell classes (all from the DCN) had too few cells for effective classification. These included 1 “Type-B” cell [30], 1 chestnut cell [31], 7 giant cells, 2 “horizontal bipolar” cells (small neurons in the pyramidal cell layer of the DCN with a bipolar morphology where the moderately spiny dendrites reside mainly within the pyramidal cell layer), 2 molecular layer stellate cells [32], 3 unipolar brush cells [31,33], and 3 cells that could not clearly be identified on comparison with the literature. These were not included in the analyses.

**Fig 1.**
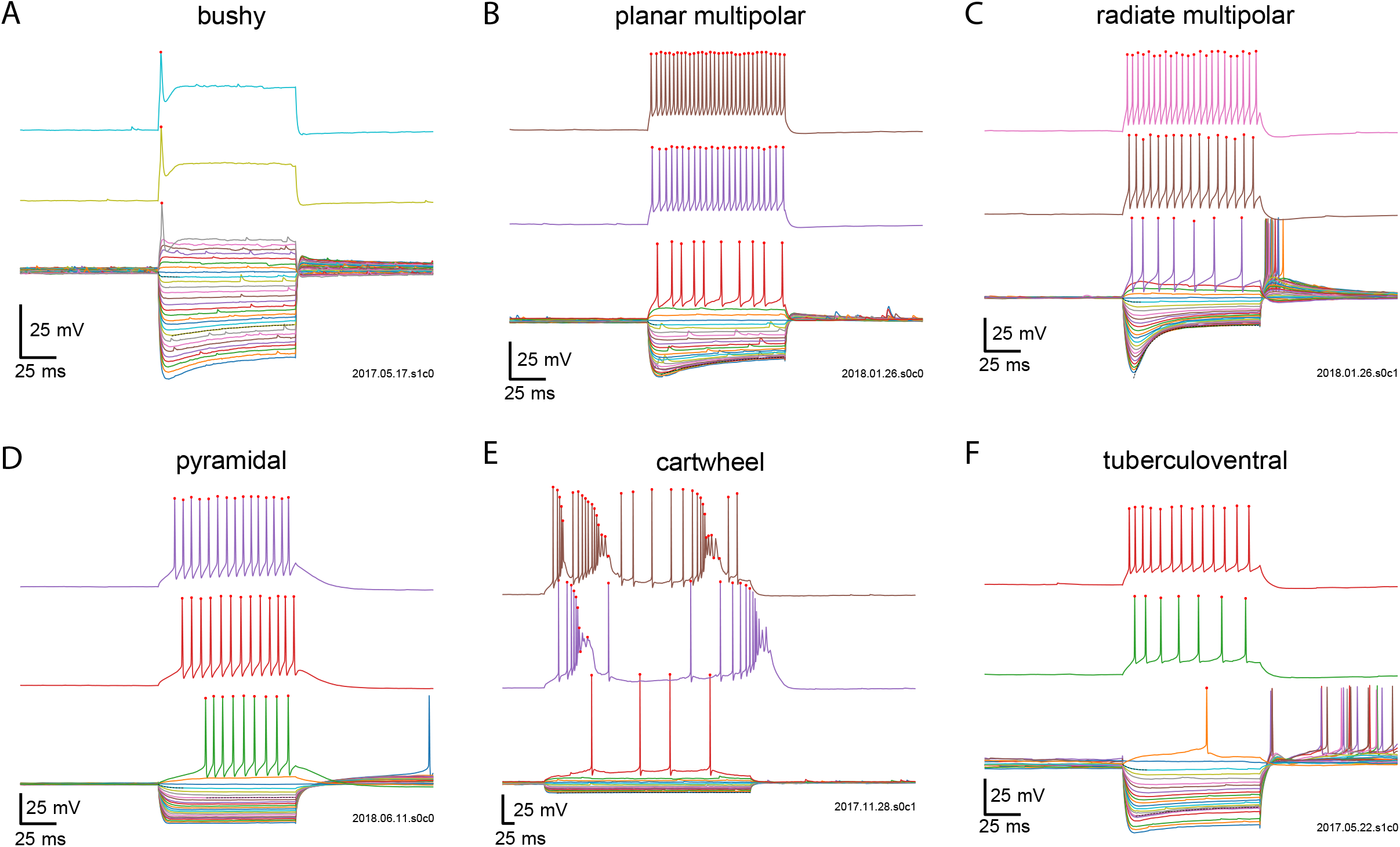
Examples of discharge patterns and passive responses, for different levels of current injection for 6 classes of cochlear nucleus neurons. A. Bushy; B. planar multipolar; C. radiate multipolar; D. pyramidal; E. cartwheel; F. tuberculoventral.. Red dots indicate spikes evoked by current injection.

In order to classify the principal cell types, we extracted a set of measurements from the current-voltage and spiking responses (N = 162 cells). These are illustrated in Figs 2 and 3. Fig 2 summarize the passive properties of the cells (columns) against the cell types (rows). From this, differences in the resting membrane potential, input resistances and time constants can be appreciated between the groups. In addition, measurements of the time constant of the hyperpolarizing sag, and a previously-used measure [23], the B/A sag ratio, also show clear differences between the cells, with neurons from the DCN generally showing weaker I_h_.

**Fig 2.**
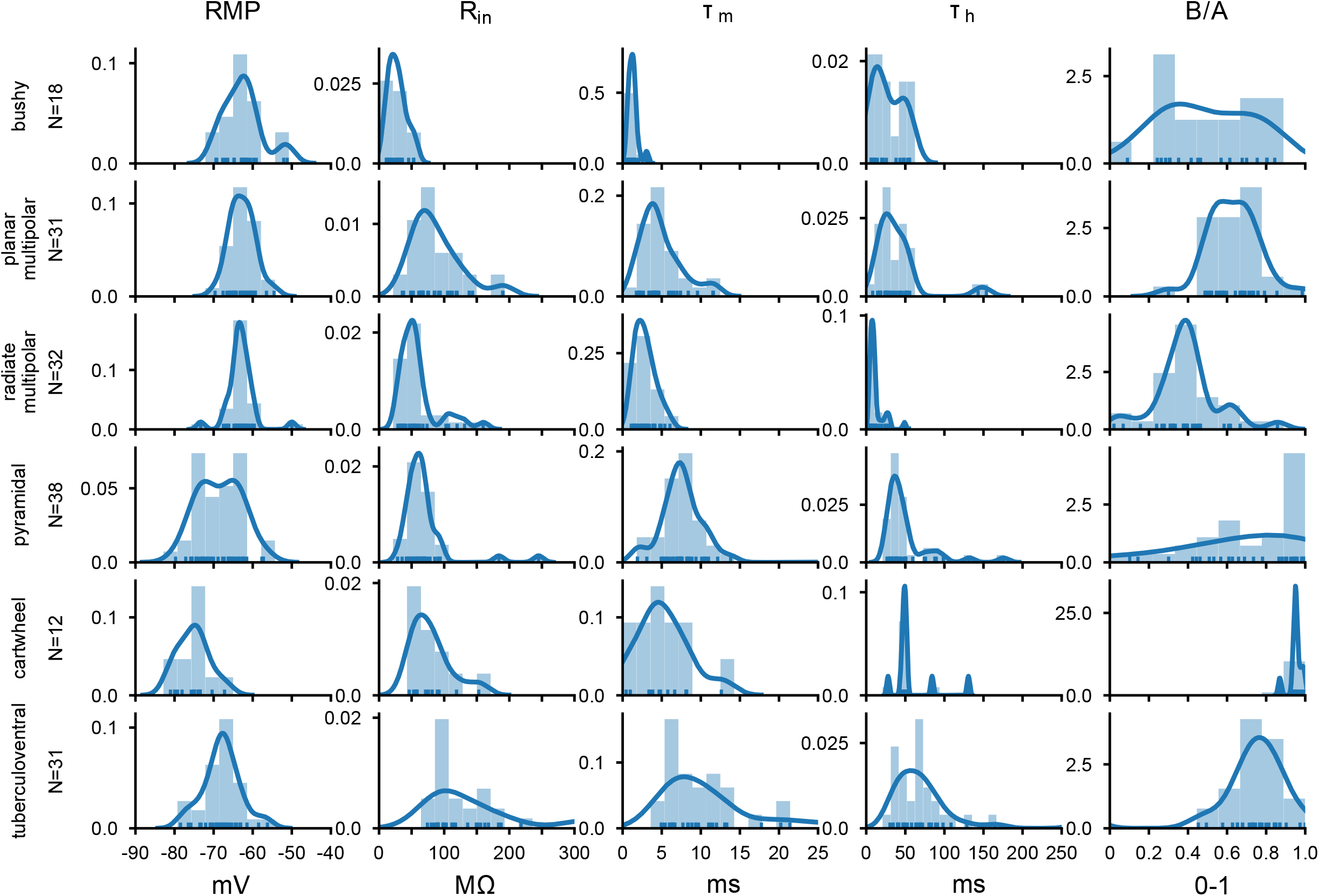
Summary of passive measurements from responses to hyperpolarizing current pulses across the population of CN neurons. RMP: resting membrane potential; R_in_: input resistance; τ_m_: membrane time constant; τ_h_: time constant of repolarizing sag from traces near −80 mV; B/A: steady-state over peak voltage deflection for hyperpolarizing sag (from Fujino and Oertel, 2001). The ordinate indicates the population density based on the kernel density estimate (blue line). The histograms shows the distribution of values from the population cells for each type and measure.

**Fig 3.**
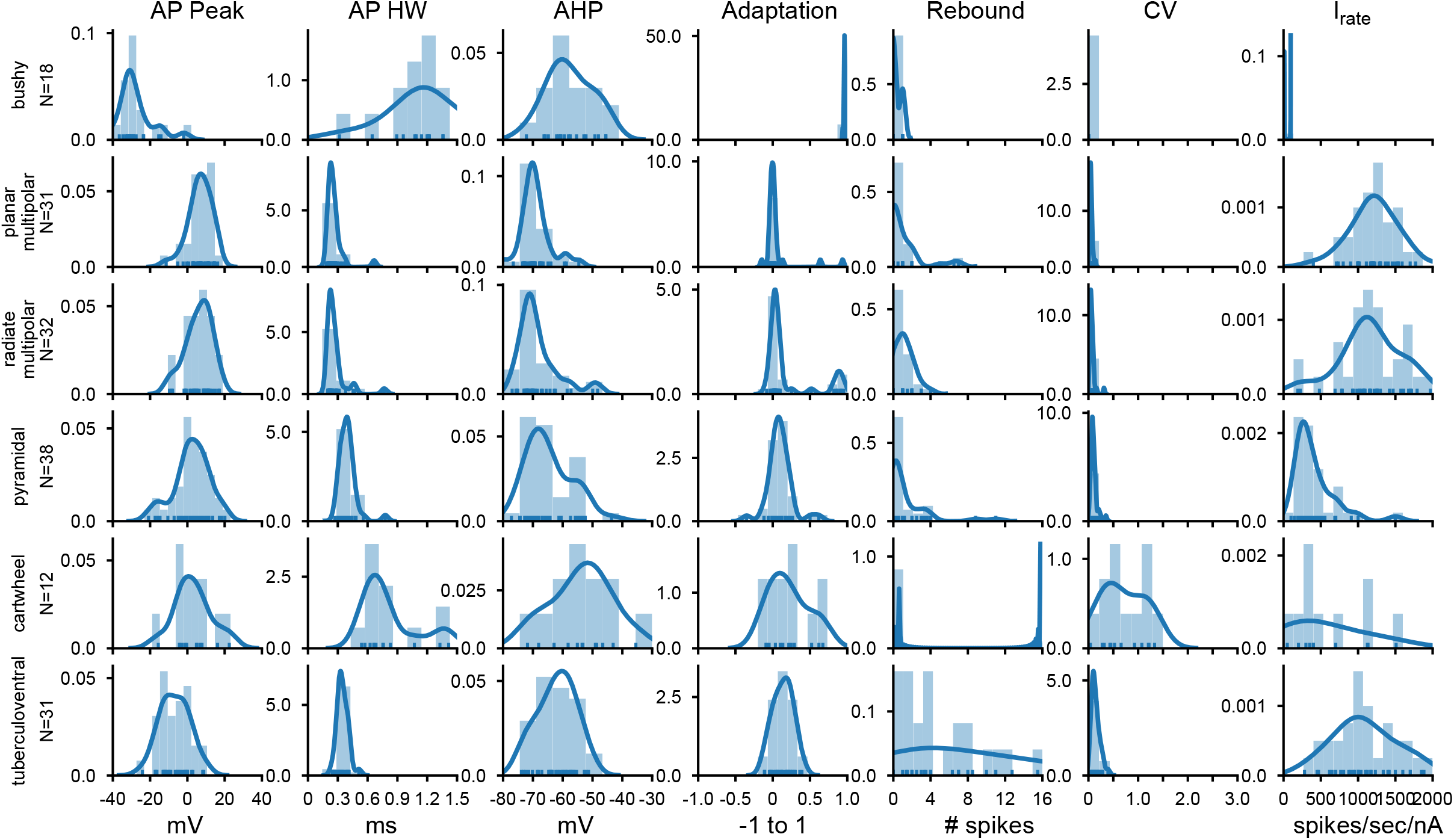
Summary of action potential shape and firing patterns across the population of CN neurons. AP Peak: action potential peak potential; APHW: action potential half-width; AHP: action potential afterhyperpolarization; Adaptation: adaptation calculated from the response to a suprathreshold current injection; Rebound: count of rebound action potential after the end of the hyperpolarizing current injection; CV: coefficient of variation of interspike intervals; Irate: slope of the current-firing relationship for current levels just above spike threshold. The ordinate indicates the population density based on the kernel density estimate (blue line). The histograms show the distribution of values from the population cells for each type and measure.

Fig 3 shows measures of action potential shape and firing properties. Again, population-based distinctions are evident, such as the relatively small and wide action potentials of bushy cells, and the tendency of tuberculoventral and some cartwheel cells to show rebound responses, and the wide coefficient of variation of firing of the cartwheel cells. The firing rate slope measured near threshold also was lowest for bushy, pyramidal and cartwheel cells, and highest for the planar and radiate multipolar cells, and tuberculoventral cells.

Next, we submitted the data to a LDA, using all of the parameters measured in Figs 2 and 3. Data were first standardized for each measure before being submitted to the LDA. The standardization rescaled the individual measurements for each measurement type so that it had a zero mean and a unit standard deviation. Fig 4 illustrates the first 3 components of the LDA, with each cell colored by its classified type, in 3 views (Fig 4A, B, C). The LDA effectively separated the different types of cells into distinct spaces. The bushy and cartwheel cells were the most separated from the remainder of the regular firing cells. Interestingly these two cell groups did not form tight clusters, suggesting some diversity in their properties. The pyramidal and tuberculoventral cells were clustered next to each other, although with minimal overlap. The radiate and planar multipolar cells formed two slightly overlapping clusters that were largely separate from all other cell classes. Note that although most of the bushy, planar and radiate multipolar cells were recorded in CBA mice, those cells recorded from the NF107::Ai32 mice (FVB and C57Bl/6 backgrounds; solid symbols) were close to the measures of the CBA populations, although they were slightly separated in one of the first 3 axes, as more clearly seen in Fig 4D, where only cells from the VCN are shown. Cross-validation of the LDA yielded an estimated accuracy of 0.79 (+/− 0.31).

**Fig 4.**
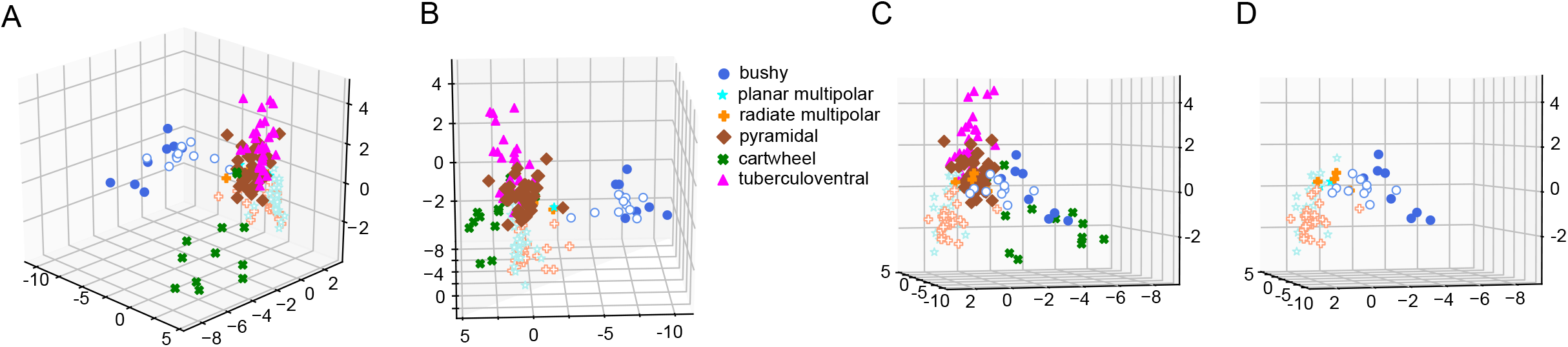
LDA with supervised clustering by cell morphology. A. The first 3 (largest) LDA components are shown in a perspective view. The axes represent the projections of each cell on to the 3 largest components from the standardized data (zero mean, unit standard deviation; therefore there are no units). Note the clear separation of the bushy cell and cartwheel cells populations from the rest of the CN neurons. Although the other populations are close together, they are also separated as can be appreciated by comparing the different views. Cells from CBA mice are shown with open symbols; cells from the NF107 mice are shown with closed symbols. B, C. Two other perspective views (rotated) of the same data as in A. D. A view of the data for the VCN cells only (bushy, planar multipolar and radiate multipolar) for clarity. This is the same perspective view as panel C.

We also submitted the data set to a standard principal components analysis, following the same standardization across cells for each measure (Fig 5). In this case, the supervisory classifier (cell morphology) was not used in the initial classification. The PCA resulted in clusters of cells from the same morphological class, but these had greater overlap than with the LDA. Cross-validation of the PCA data yielded a low accuracy of 0.17 (+/− 0.086).

**Fig 5.**
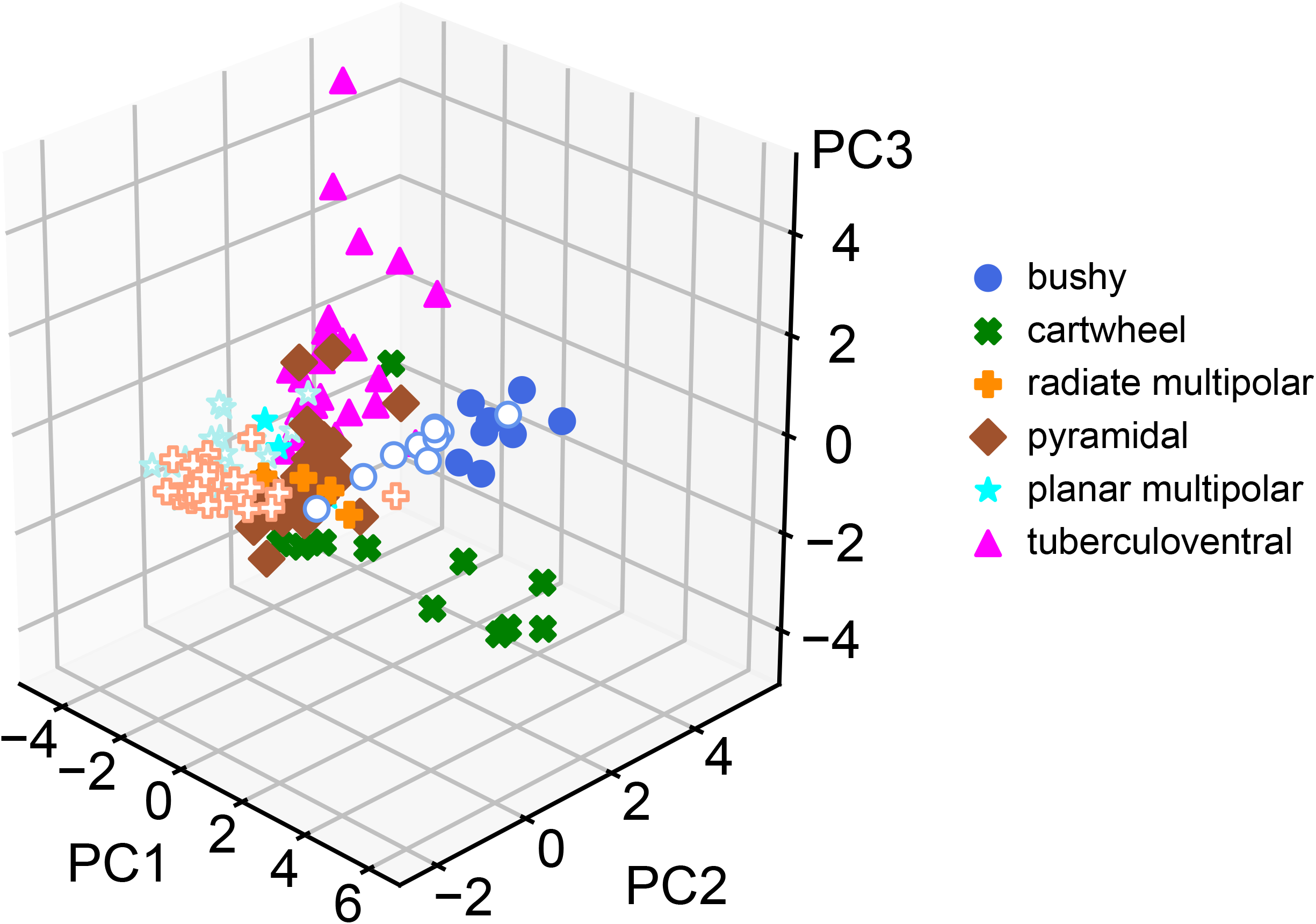
Principal components analysis (PCA) on the same data set as Fig 4. The PCA method is not supervised by cell type, and so the results depend only on factors that maximize the variance. The PCA successfully separated the cell classes, but did not have good accuracy with cross-validation. The view shown is the same format as Fig 4A.

In order to determine which measures provided the most information in the LDA classification, we performed the LDA using combinations of measures, from individual measures through all available measures, and estimated the accuracy of the classification across all cells by dividing the data into training and testing sets. The accuracy as a function of the number of combined measures is shown in Fig 6. As expected, the accuracy improves as new measures are added, up until about 6 parameters, at which point the accuracy plateaus. However, the overall *worst* accuracy continues to improve as more measures are added. The black line indicates the mean of the best 5 combinations of measures. From this we conclude that some of the measures are possibly redundant and that some measures may be non-informative. With 7 or 8 combined measures (where the largest number of combinations was tested), AP height, R_in_, RMP, τ_h_ and I_rate_ occurred together in each of the 5 most accurate runs. Similar, but not identical distributions were present for 8 combined measures. Note that the accuracy of each point includes a standard deviation estimate (not shown) as it is the result of multiple runs with different subsamples of training and test cells, so the best measures can vary with an arbitrary threshold, and there is no single “optimal” set. With the number of cells in the sample and the large number of parameters, the SD can be 15-20% of the mean value.

**Fig 6.**
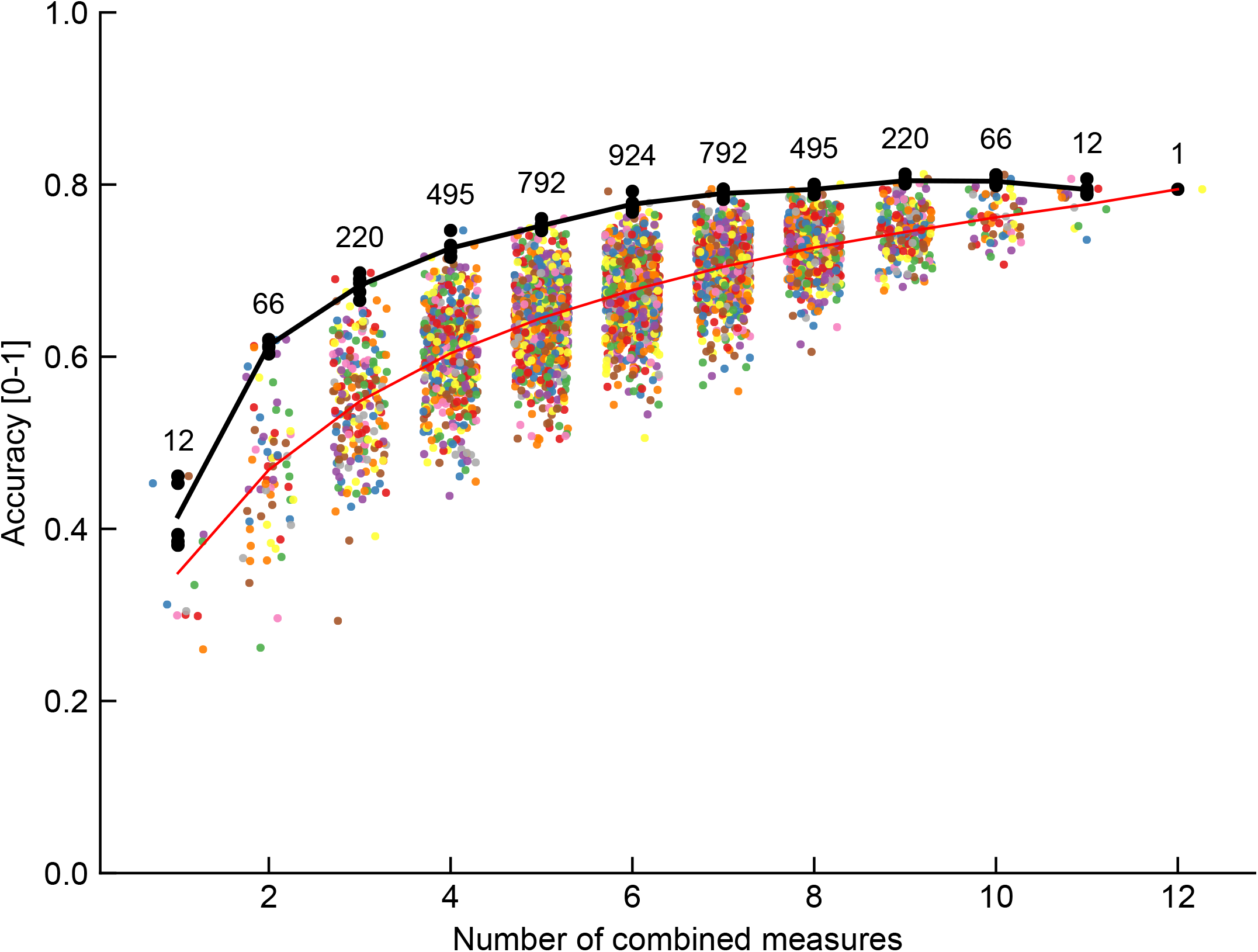
Accuracy of separation for different number of combined measures, using the LDA separation score for each set of measures. All 12 measures were considered in all possible combinations in groups from 1 to 12 (the number of combinations are shown above the data), and each combination is plotted as a point along the ordinate representing the number of combinations. The mean score is indicated by the red line. The average of the best 5 scores for each combination set are plotted as a black line. In general, including more measures improves the accuracy of the separation of groups.

To further investigate those factors that drove the prediction accuracy, we performed the same analysis using the R package flipMultivariates. Table 1 summarizes the prediction accuracy by cell type, and provides mean measures for each of the parameters. Although all parameters provide a significant contribution (r^2^) to the separation, the five that accounted for the largest proportions of the variance (r^2^ > 0.50) were the AP height, AP half width, the adaptation measure, the coefficient of variation of interspike intervals, and the firing rate slope (I_rate_). However, all of the measures showed a significant contribution.

**Table 1.**
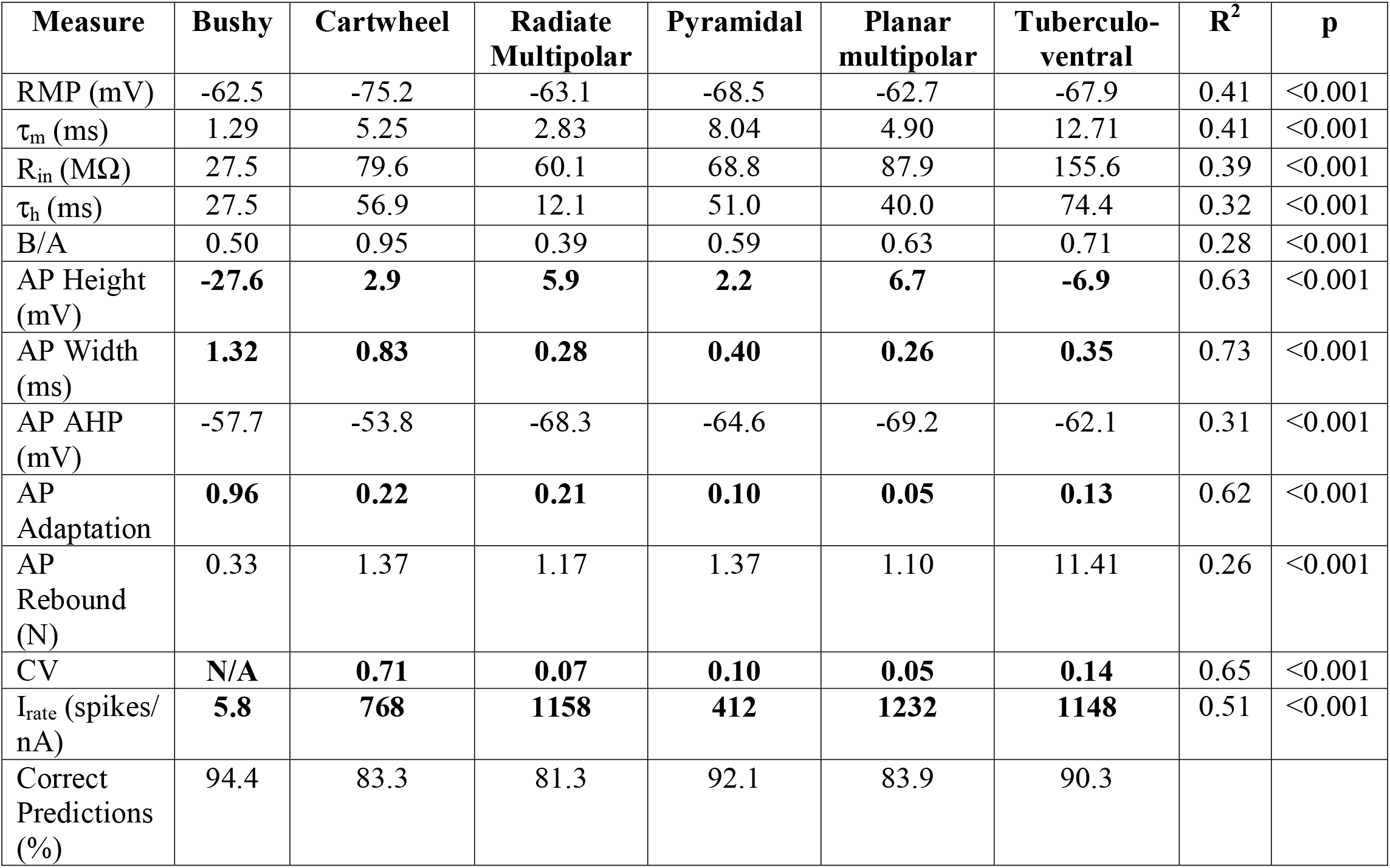
Relations between the predictors and each cell class, indicating the number of correct predictions overall, and for each class. The top predicting variabiles (largest R^2^ values) are highlighted in bold. N = 162 cases used in estimation. Null hypotheses: two-sided; multiple comparisons correction: False Discovery Rate correction applied simultaneously to entire table. N/A: not applicable. See Methods and Materials for details of measurements.

Fig 7 plots the overall prediction accuracy against the observed classifiers. Although the overall accuracy was fairly high (87.1%), there were several confounds. The most common of these was radiate vs. planar multipolar, which occurred in about 25% of the cells in these groups. The next most common confound was misclassifying planar multipolar and tuberculoventral cells as pyramidal cells, followed by pyramidal cells being misclassified as tuberculoventral and planar multipolar cells. As these cells all fire regularly, and have similar measures on other properties, this is not surprising.

**Fig 7.**
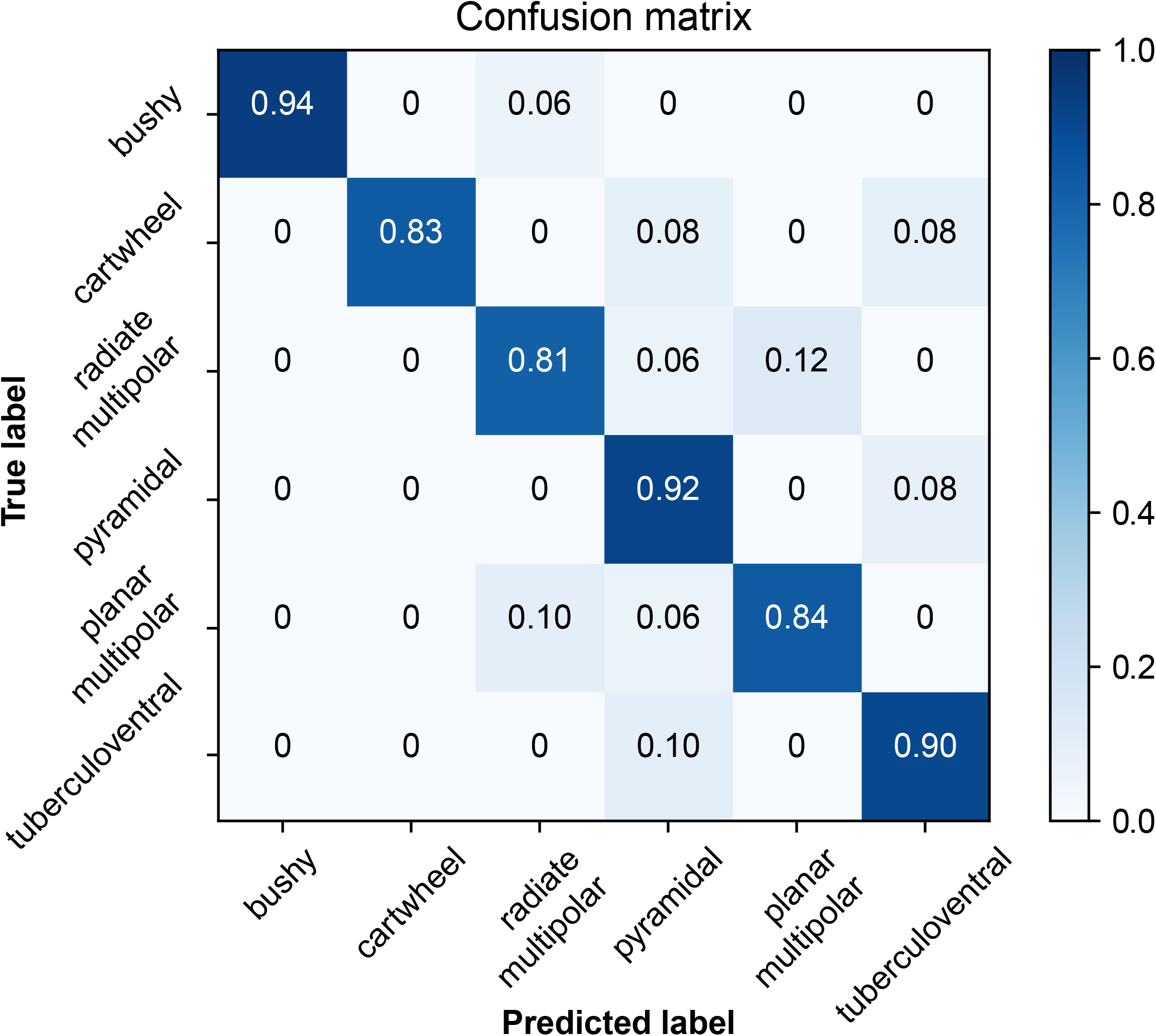
Prediction accuracy table (confusion matrix) by cell type. The most frequent mis-classifications occurred amongst cells with similar general firing patterns, such as planar and radiate multipolar cells, and pyramidal and tuberculoventral cells. This table was generated in R using data from all 12 measures. The numbers in each box indicate the proportion of correct classifications for each cell type.

## Discussion

We find that cells thoughout the cochlear nucleus can be classified by firing patterns, action potential shapes, and responses to hyperpolarizing steps, as well as their morphology. Although not surprising given the long history of the identification of various electrophysiological features as identifying characteristics of different morphological cell classes, the extension of these comparisons across classes that fire regularly, and the consistency of these measures in a population of cells taken from adult mice extends the previous observations made in younger mice. In addition, these results suggest that cells can be reasonably classified, at least coarsely, according to their electrophysiological signatures.

We found that the linear discriminant analysis can be used to classify cells based on 12 measured electrophysiological parameters with ~80% success, although there was little improvement when using more than 7 parameters. This requires that the LDA coefficients be estimated from a standardized (training) set of cells before application to an unidentified population. The selection of the training cell, and the quality of that data set, is critical to the success of the technique. Cell morphology should be positively identified, and ambiguous cases discarded, although it is essential to include a full range of cell properties in each class. The measurements should be complete and precise for each cell. Our data set has three limitations in part because it was collected as part of a different experiments. The first is that due to different levels of hyperpolarizing current injection, the voltages reached with hyperpolarizing pulses were not consistent across cells, so that the estimates of the magnitude of the hyperpolarizing sag are influenced by the variability of the voltages reached. The second is that for depolarizing pulses, not all cells reached saturation of their firing rates. For this reason, we did not include maximal firing rates as a measure, but rather focused on the discharge patterns closer to spike threshold. A third limitation is that the current steps near spike threshold were, in some cells, too coarse to precisely define a threshold current. These limitations partly reflect the different input resistances of neurons, as an example, hyperpolarizing pulses strong enough to damage tuberculoventral cells often fail to hyperpolarize pyramidal cells to −80 mV.

Classification errors principally occurred between cell classes with similar firing patterns, such as planar and radiate multipolar cells, and between pyramidal and tuberculoventral cells. In addition, the TV and bushy cells show significant dispersion in the first 3 dimensions of the LDA. This may indicate variability in the intrinsic excitability of these cell classes as noted before [12,34,35], or possibly the existence of distinct subclasses. This dispersion was also evident in the unsupervised PCA analysis. Substantially larger datasets of identified cells from individual strains would be needed to clarify the existence of such subclasses in the electrophysiology. An improvement in the classification would be expected to result from the inclusion of additional parameters such as maximal firing rates and the time courses of synaptic events. In addition, in our data set there was some overlap between the planar and radiate multipolar cells. This may in part reflect the limitations of classification for these two similarly firing cell types, but may also indicate the limitations of our morphological classification method, which was qualitative and relied on fluorescent images of the cells collected during the experiments. The qualitative morphological classification of DCN neurons is much easier, so the overlap between the pyramidal and tuberculoventral cells probably represents the limitations of the measurement parameters used, although it could also reflect a true confluence of intrinsic excitability.

Part of the dispersion in the VCN cell classes may reflect strain or preparation differences, as the cells from CBA mice were slightly offset from those from the NF107::Ai32 mice along the second LDA component. The strain difference is reminiscent of the differences in HCN channels seen in bushy, planar multipolar and octopus cells between ICR and a knockout on a hybrid 129S and C57Bl/6 background [3]. This raises a cautionary flag that the LDA should be trained on data acquired from cells recorded from animals of the same genetic background (and age and preparation techniques) if it is to be used to categorize cells from a novel data set.

## Acknowledgements

This research was supported by the National Institute on Deafness and other Communicative Disorders of the National Institutes of Health grants R01 DC004551 to PBM and R03 DC013396 to RX

## Notes

#### Summary of Updates

Minor corrections to # of cells, updated text to reflect re-analysis, Figures 2-7 and table 1 revised to be consistent with updated analysis. The differences are minor and quantitative and do not affect any conclusions or statistics.

